# Brain age prediction in schizophrenia: does the choice of machine learning algorithm matter?

**DOI:** 10.1101/2020.07.28.224931

**Authors:** Won Hee Lee, Mathilde Antoniades, Hugo G Schnack, Rene S. Kahn, Sophia Frangou

## Abstract

**Background:** Schizophrenia has been associated with lifelong deviations in the normative trajectories of brain structure. These deviations can be captured using the brain-predicted age difference (brainPAD), which is the difference between the biological age of an individual’s brain, as inferred from neuroimaging data, and their chronological age. Various machine learning algorithms are currently used for this purpose but their comparative performance has yet to be systematically evaluated.

**Methods:** Six linear regression algorithms, ordinary least squares (OLS) regression, ridge regression, least absolute shrinkage and selection operator (Lasso) regression, elastic-net regression, linear support vector regression (SVR), and relevance vector regression (RVR), were applied to brain structural data acquired on the same 3T scanner using identical sequences from patients with schizophrenia (n=90) and healthy individuals (n=200). The performance of each algorithm was quantified by the mean absolute error (MAE) and the correlation (R) between predicted brain-age and chronological age. The inter-algorithm similarity in predicted brain-age, brain regional regression weights and brainPAD were compared using correlation analyses and hierarchical clustering.

**Results:** In patients with schizophrenia, ridge regression, Lasso regression, elastic-net regression, and RVR performed very similarly and showed a high degree of correlation in predicted brain-age (R>0.94) and brain regional regression weights (R>0.66). By contrast, OLS regression, which was the only algorithm without a penalty term, performed markedly worse and showed a lower similarity with the other algorithms. The mean brainPAD was higher in patients than in healthy individuals but varied by algorithm from 3.8 to 5.2 years although all analyses were performed on the same dataset.

**Conclusions:** Linear machine learning algorithms, with the exception of OLS regression, have comparable performance for age prediction on the basis of a combination of cortical and subcortical structural measures. However, algorithm choice introduced variation in brainPAD estimation, and therefore represents an important source of inter-study variability.

## Introduction

Schizophrenia is a major mental illness that presents with positive and negative symptoms^1^ and brain structural abnormalities.^2-7^ Lower intracranial volume is a consistent finding that predates illness onset and reflects early developmental vulnerability to the disorder.^8, 9^ Further abnormalities involve widespread reductions in cortical thickness and subcortical volumes^2-7^ which generally show a steeper age-related decline than that observed in healthy individuals.^2, 10-13^ Collectively, these findings suggest that schizophrenia is associated with a lifelong deviation from normative trajectories of brain organization.

Application of machine learning algorithms to neuroimaging data offers a new lens for the examination of brain structural deviation in schizophrenia. Using these algorithms, it is possible to generate estimates of the biological age of an individual’s brain (i.e., brain-age) by comparing their neuroimaging data against a normative population dataset of the same neuroimaging features.^14^ In each individual, subtracting their chronological age from the neuroimaging predicted brain-age generates a “brain-predicted age difference” (brainPAD) score, also referred to as brain age gap estimation (BrainAGE)^15, 16^; a positive score indicates that the brain-age of an individual is “older” than their actual age, and a negative score reflects the inverse.

A recent study using brain structural data from 35,474 healthy individuals and 1,110 patients with schizophrenia demonstrated that schizophrenia is associated with an increase in mean brainPAD of moderate effect size (Cohen’s *d* effect size = 0.51).^17^ Five further brain structural studies have focused on estimating the years of brain-age acceleration in schizophrenia using various machine learning algorithms.^16, 18-21^ Koutsouleris and colleagues used a support vector regression (SVR) algorithm trained on data from 800 healthy individuals, and found that patients with schizophrenia (n=141) had a mean brainPAD of 5.5 years.^18^ Schnack and colleagues also used SVR trained on a sample of 386 healthy individuals, and showed that discovery (n=341) and validation sample (n=60) of patients with schizophrenia had a mean brainPAD of 3.8 years and 5.6 years, respectively.^16^ Using relevance vector regression (RVR) trained on data from 70 healthy individuals, Nenadic and colleagues reported that patients with schizophrenia (n=45) had a mean brainPAD of 2.6 years.^19^ A similar mean brainPAD of 2.6 years in patients with schizophrenia (n=43) was reported by Hajek and colleagues using RVR trained on data from 504 healthy individuals.^20^ Using random forest regression trained on data from 50 healthy individuals, Shahab and colleagues showed that patients with schizophrenia (n=81) had a mean brainPAD of 7.8 years.^21^ Other regression algorithms commonly used to estimate brain-age in neuropsychiatric conditions include ordinary least squares (OLS) regression,^22, 23^ least absolute shrinkage and selection operator regression (Lasso),^24, 25^ ridge regression,^26, 27^ and elastic-net regression.^28^

At present, it is unclear whether the performance of these machine learning approaches is comparable when estimating brainPAD. In principle, brainPAD can be influenced by the specific characteristics of the study sample, the choice of neuroimaging features entered into the machine learning models or the type of the machine learning algorithm used. In the present study, we focus specifically on the comparative performance of the commonly used machine learning algorithms in estimating brainPAD in schizophrenia. To achieve this, we leveraged the availability of brain structural magnetic resonance imaging (MRI) data from 200 healthy individuals and 90 patients with schizophrenia acquired using identical scan sequences on the same 3T scanner and processed using the same image processing pipelines. We focused exclusively on six linear machine learning algorithms (OLS regression, ridge regression, Lasso regression, elastic-net regression, SVR, and RVR) due to their interpretability and resilience to over-fitting.^29^ The brain features entered in each algorithm were regional measures of cortical thickness, cortical surface area and subcortical volume derived using FreeSurfer-implemented parcellation and segmentation of each individual’s imaging dataset. We chose these measures as they are amongst the most commonly extracted imaging features in brain imaging studies of schizophrenia^3, 4^ and have high translational potential.

## Materials and Methods

### Sample

The patient sample was recruited via clinician referrals from the psychiatric services of the Mount Sinai Health System at the Icahn School of Medicine at Mount Sinai (ISMMS), New York. All patients were screened to ensure that they met diagnostic criteria of schizophrenia according to the Diagnostic and Statistical Manual of Mental disorders, Fifth Edition (DSM-5).^1^ Healthy individuals were recruited from the same local area via advertisements and did not have a personal or family history of major psychiatric disorders. Additionally, all participants were screened to exclude individuals with IQ<70, medical or neurological disorders including head trauma or loss of consciousness, lifetime history of DSM-5 substance use disorder, and contraindications to MRI. The study was approved by the Institutional Review Board of the Icahn School of Medicine at Mount Sinai and all participants gave written informed consent.

Diagnostic status was established in all participants using the Structured Clinical Interview for DSM-5, research edition.^30^ The Brief Psychotic Rating Scale (BPRS)^31^ was used to evaluate symptoms in patients; this instrument yields subscale scores for positive symptoms, negative symptoms, anxiety/depression and agitation/disorganization^32^ (Supplementary Table S1). Additional information was collected on age of onset and duration of illness in patients. All patients were scanned while medication-free for a minimum of two weeks.

The characteristics of the patient sample are shown in Table 1. The median duration of illness was 4 years with an interquartile range of 1-7 years. To ensure robust calculation of age prediction from the neuroimaging data, we used structural magnetic resonance imaging (MRI) scans from 200 healthy individuals (110 men and 90 women) aged 19 to 70 years (mean age = 37 years, standard deviation = 12.4 years).

**Table 1.** Sample characteristics.

| **Table 1. Sample characteristics** |  |  |
| --- | --- | --- |
|  | **Patients with**<br>**schizophrenia (N=90)** | **Healthy individuals**<br>**(N=200)** |
| **Age (years)** | 27.43 (7.51) | 36.96 (12.39) |
| **Age range (years)** | 18 - 52 | 19-70 |
| **Sex (male/female)** | 69/21 | 110/90 |
| **Age of onset (years)** | 21.71 (5.46) | n/a |
| **BPRS: positive subscale score** | 11.18 (5.39) | 4.00 (0) |
| **BPRS: negative subscale score** | 6.97 (3.77) | 3.02 (0.14) |
| **BPRS: anxiety/depression subscale**<br>**score** | 8.83 (5.28) | 4.02 (0.14) |
| **BPRS: agitation/disorganization**<br>**subscale score** | 8.35 (5.40) | 6.02 (0.14) |
| Continuous variables are shown as mean (standard deviation); BPRS=Brief Psychiatric Rating Scale;<br>Definition of the BRPS subscale scores in Supplementary Table S1; n/a=not applicable. |  |  |

### Neuroimaging data acquisition and processing

All participants were scanned on the same 3T Skyra scanner (Siemens Medical Systems, Erlangen, Germany) at ISMMS using identical acquisition protocols. Details of the neuroimaging data acquisition, preprocessing, quality assurance and imaging measures extraction are provided in Supplementary Material. Briefly, T_1_-weighted images were processed using FreeSurfer 6.0 (http://surfer.nmr.mgh.harvard.edu). The Desikan-Killiany atlas^33, 34^ was used to parcellate the brain into cortical regions while subcortical segmentation was performed using the probabilistic atlas in FreeSurfer.^35^ This procedure generated measures of total intracranial volume and regional measures of cortical thickness (n=68), cortical surface area (n=68) and subcortical volumes (n=16) (Supplementary Table S2).

### Machine learning algorithms

We used a linear regression model to predict brain-age using brain regional structural measures. The linear model can be formalized as follows:

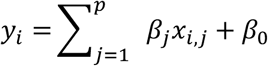

where *y*_*i*_ is the age of the *i*^*th*^ individual, *p* is the number of features, *x*_*ij*_ is the value of *j*^*th*^ feature of the *i*^*th*^ subject, and *β*_*j*_ is the regression coefficient. We evaluated the following six algorithms:

#### (1) Ordinary least squares (OLS) regression

OLS regression algorithm fits a linear model by minimizing the residual sum of squares between the observed *y*_*i*_ in the training dataset (*i*=1,..,*N*, the sample size) and the values *f*(*x*_*i*_) predicted by the linear model. The object function is as follows:

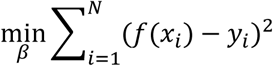

where *y*_*i*_ is the actual value of the chronological age. The least squares solution was computed using the singular value decomposition (SVD).

#### (2) Ridge regression

Ridge regression is a regularized linear model that minimizes the sum of the squared prediction error in the training data and an L2-norm regularization, i.e., the sum of the squares of regression coefficients.^36^ The object function is as follows:

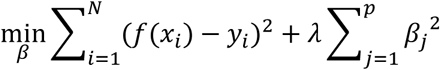

The tuning parameter *λ* controls the model’s complexity. If *λ*= 0, ridge regression becomes a traditional linear regression model. The optimal choice of *λ* parameter in this study was based on 10-fold cross-validation (see below).

#### (3) Least absolute shrinkage and selection operator (Lasso) regression

Lasso is another form of regularized linear regression using an L1-norm penalty, aiming to minimize the sum of the absolute value of the regression coefficients.^37^ The objective function is as follows:

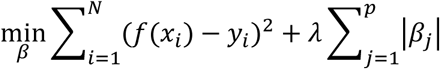

The L1-norm regularization tends to set most coefficients to zero and retains one random feature among the correlated ones, thus resulting in a sparse predictive model that facilitates optimization of the predictors and reduces the model complexity.

#### (4) Elastic-net regression

This linear regression model combines both L1-norm (i.e., Lasso regression) and L2-norm (i.e., ridge regression) regularizations in the OLS loss function.^38^ The object function is as follows:

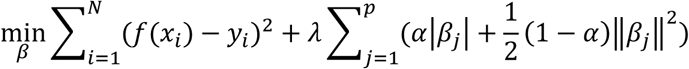

This allows the number of selected features to be larger than the sample size while achieving a sparse model. A hyperparameter parameter *α* (between 0 and 1) is used to control the relative weighting of the L1-norm and L2-norm contributions. The optimal choice of *α* parameter in this study was based on 10-fold cross-validation (see below).

#### (5) Linear support vector regression (SVR)

Linear SVR aims to find a function *f*(*x*_*i*_ ) whose predictive value deviates by no more than a required accuracy *ε* from the actual *y*_*i*_ for all the training data while maximizing the flatness of the function.^39^ Flatness maximization is implemented using the L2-norm regularization by minimizing the squared sum of the regression coefficients. The object function is as follows:

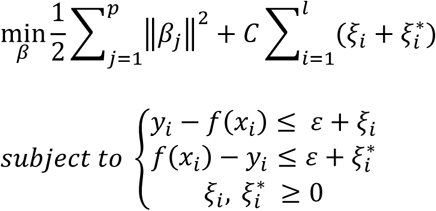

where *l* is the quantity of ‘support vectors’, which are the samples that deviate by more than *ε* from the actual *y*_*i*_ used to fit the model. A parameter *C* regulates the smoothness of function *f*(*x*_*i*_ ). The optimal choice of *C* parameter in this study was based on 10-fold cross-validation (see below).

#### (6) Relevance vector regression (RVR)

RVR is a Bayesian sparse learning model and has an identical functional form to SVR.^40^ The function is as follows:

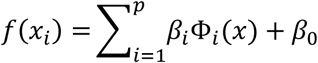

where *β* = (*β*_0_,…, *β*_*p*_) is a vector of weights and Ф_*i*_(*x*) = *K*(*x,x*_*i*_) is a linear kernel function defining the basis function. The sparsity of RVR is induced by the hyperpriors on model parameters in a Bayesian framework with the maximum a posteriori (MAP) principle. RVR determines the relationship between the target output and the covariates by enforcing sparsity. The L1-norm-like regularization used in RVR encourages the sum of absolute values to be small, which often drives many parameters to zero and provides significantly few basis functions. Notably, RVR has no algorithm-specific parameter.^40^

The scikit-learn library (version 0.23.1) was used to implement OLS regression, ridge regression, Lasso regression, and elastic-net regression (http://scikit-learn.org/),^41^ the LIBSVM function in MATLAB (Mathworks, Natick, MA) was used to implement SVR (https://www.csie.ntu.edu.tw/~cjlin/libsvm/),^42^ and the PRoNTo toolbox (http://www.mlnl.cs.ucl.ac.uk/pronto/) was used to implement RVR.^43^

### Brain-age prediction framework

The six regression algorithms were applied separately to the data but the analytic procedure was common to all and involved the following steps: (i) prior to modeling, each neuroimaging measure was linearly scaled so that all values in the feature set ranged between 0 and 1; (ii) a nested 10-fold cross-validation (10F-CV) was applied to the dataset of healthy individuals which was randomly split into 10 equal-sized subsets. For each cross-validation, one subset was left out as the test subset while the remaining nine subsets were used together as the training set for estimating the model parameters. These parameters were then applied to the left-out subset. Specifically, for ridge regression, Lasso regression, elastic-net regression, and SVR, cross-validation procedure was applied with an outer 10F-CV to evaluate model generalizability and an inner 10F-CV to determine the optimal parameters (λ, α, or C) for these algorithms; (iii) the performance of each algorithm was quantified by the mean absolute error (MAE) and the correlation (R) between predicted ‘brain-age’ and chronological age, respectively, averaged across all cross-validation folds; (iv) the performance of each algorithm was first evaluated in the sample of healthy individuals with respect to different types of regional brain structural measures alone (regional measures of cortical thickness and surface area and subcortical volumes) and in combinations. As shown in Supplementary Table S3, all models performed better when all brain structural features were included in the model; therefore, from here onwards we examined only these models with all combined structural features; (v) for each algorithm, the regression weights for each brain region, as derived from healthy individuals, were recorded and used for the comparative evaluation of the algorithms; the absolute value of these weights represents the importance of the corresponding features in the brain-age prediction of the model^44^; (vi) the regional regression weights of each algorithm were then applied to the brain structural data of patients with schizophrenia to derive the predicted brain-age for each patient according to each algorithm; (vii) the brain-predicted age difference (brainPAD) was calculated for each algorithm by subtracting the chronological age of each individual from their brain-age as predicted from that algorithm.

### Statistical analysis

The comparative evaluation of the algorithms was based on their similarity in predicted brain-age and brain regional regression weights. We assessed the similarity among the six algorithms using correlation analyses and hierarchical clustering with Ward’s minimum variance method for Euclidian distances implemented in MATLAB (Mathworks, Natick, MA) (Supplementary Material). Case-control differences in brainPAD were examined using the t-statistic and effect sizes were expressed in terms of Cohen’s *d*. In patients with schizophrenia, sex differences in brainPAD were tested using the t-statistic. The association between brainPAD and BPRS subscale scores was assessed using the Pearson correlation coefficient after the brainPAD values were corrected for age using a linear model to remove age-related bias.^45^ P-values were corrected using the Benjamini-Hochberg false discovery rate (FDR, *q* < 0.05) correction procedure.^46^

## Results

### Predicted brain-age

After optimizing the parameters of each algorithm in healthy individuals, the optimal models were then applied to the same structural feature set (i.e., combined intracranial volume with all the regional cortical thickness, cortical surface area and subcortical volume measures) of patients with schizophrenia. The OLS regression underperformed (MAE=12.9; R=0.10) compared to the other five algorithms which had nearly identical performance (MAE range: 7.5- 7.9; R range: 0.49-0.51) (Supplementary Table S4; Supplementary Figure S3). Individual predicted brain-ages showed high between-algorithm correlations (R > 0.94) with the exception of OLS regression which showed numerically lower correlations with all the other algorithms (R < 0.53) (Figure 1A). Hierarchical clustering of the individual predicted brain-ages in patients with schizophrenia showed that the elastic-net regression and ridge regression together formed one cluster (showing high within-cluster similarity but relatively low similarity with algorithms outside the cluster), and Lasso regression, SVR and RVR formed another cluster; by contrast, OLS regression showed a very low degree of similarity with all the other algorithms (Figure 1B).

**Figure 1.**
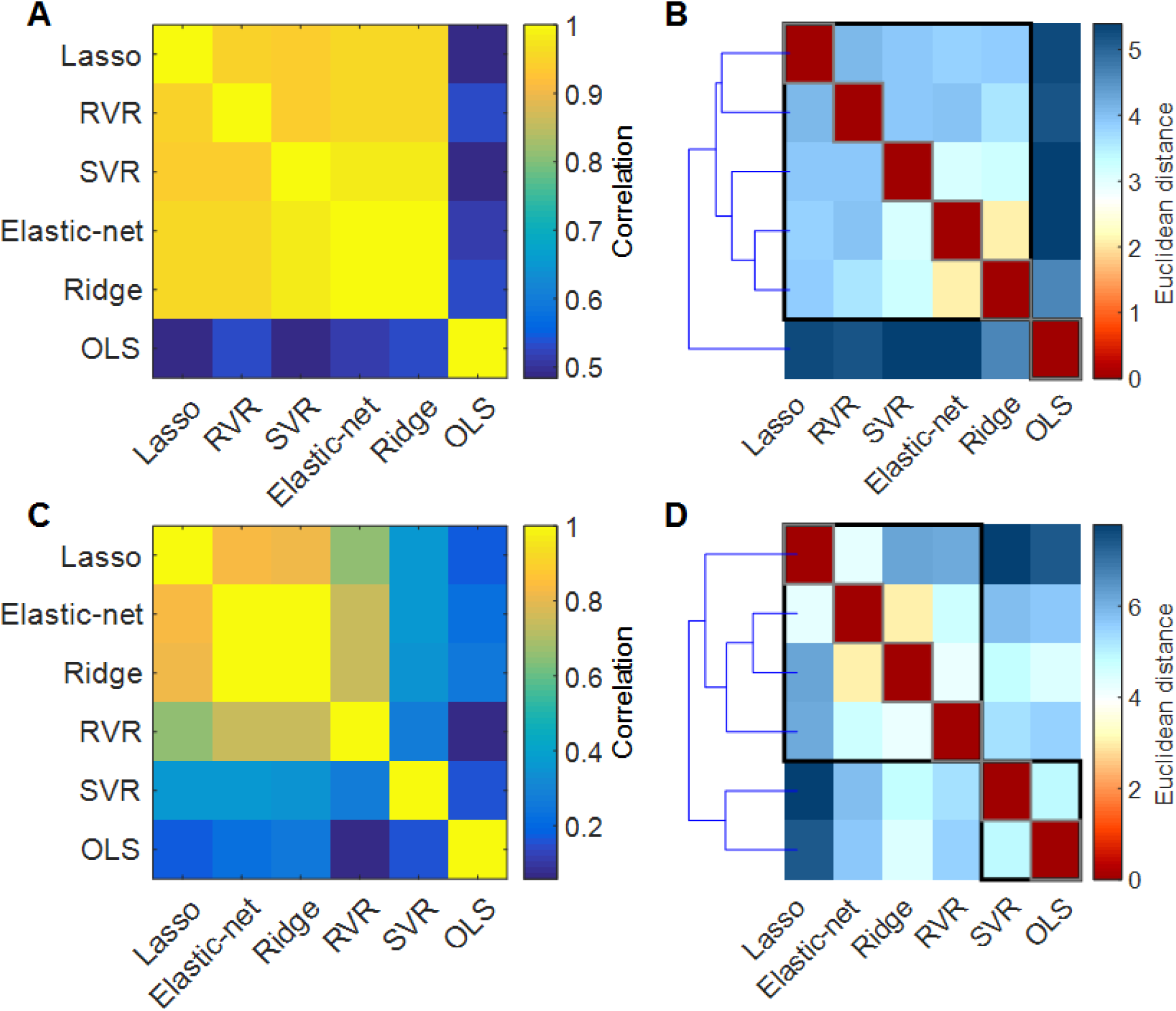
Comparative results in brain-age and brain regional regression weights from each algorithm in patients with schizophrenia. **Brain-Age:** (A) Similarity matrix representing between-algorithm correlations of individual predicted brain-age. (B) Distance matrix and dendrogram resulting from hierarchical clustering of the individual brain-age results of the six algorithms. **Brain regional regression weights**: (C) Similarity matrix representing between-algorithm correlations of the absolute regression weights of the structural features. (D) Distance matrix and dendrogram resulting from hierarchical clustering of the six algorithms in terms of brain regional regression weights. OLS: Ordinary least squares regression; Lasso: Least absolute shrinkage and selection operator regression; SVR: Support vector regression; RVR: Relevance vector regression.

### Brain regional regression weights

The regression weights for each of the 153 brain structural measures contributing to brain-age prediction in patients with schizophrenia for each of the six algorithms are shown in Supplementary Table S5 and Supplementary Figures S4-S6. Between-algorithm correlations ranged from 0.06 to 0.99 (Figure 1C). Hierarchical clustering of the absolute values of regression weights showed that ridge regression and elastic-net regression were most similar, while the results of OLS regression were the least similar to those of all other algorithms (Figure 1D).

### Brain-predicted age difference (BrainPAD)

With the exception of OLS regression, the mean brainPAD values obtained by all other algorithms were statistically higher in patients with schizophrenia compared to healthy individuals (all P_FDR_<0.001), with a range of 3.80 to 5.23 years (Supplementary Table S6; Figure 2). The effect size of case-control differences was moderate (*d* range: 0.43-0.66) (Supplementary Table S6). No sex differences in brainPAD were observed in patients with schizophrenia (P_FDR_>0.05) (Supplementary Figure S7). Positive correlations were noted between the BPRS subscale scores for positive symptoms and agitation/disorganization and the brainPAD scores for all algorithms except OLS regression (Supplementary Table S7).

**Figure 2.**
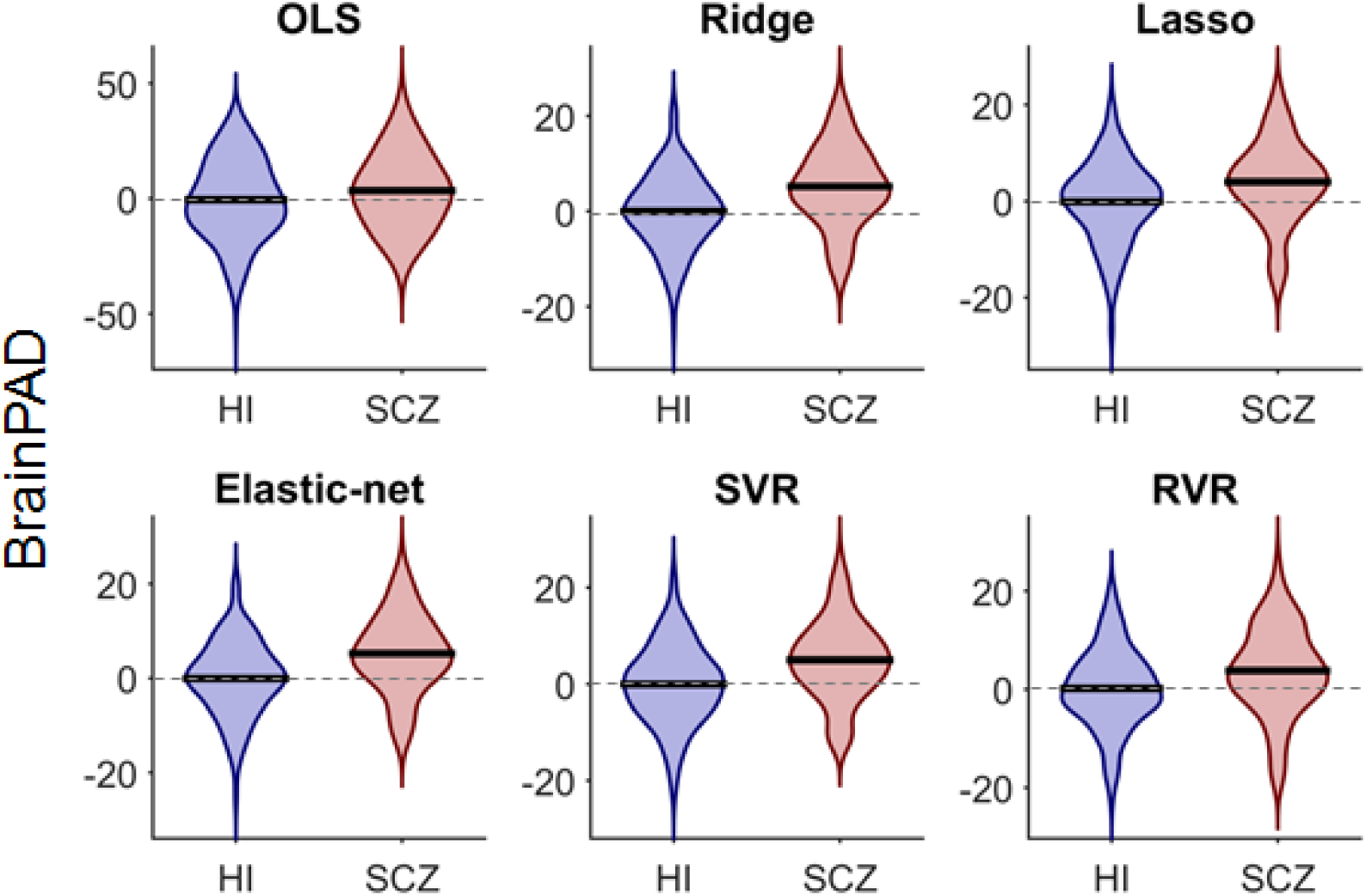
Comparative results in brain-predicted age difference (brainPAD) in patients with schizophrenia (SCZ) and healthy individuals (HI) for each algorithm. Violin plots showing the mean (black line) and distribution of individual brainPAD scores in the two diagnostic groups. With the exception of OLS regression, all other algorithms yielded significantly higher mean brainPAD (P_FDR_ < 0.001) in patients with schizophrenia than healthy individuals; Further details are provided in Supplementary Table S6. Dashed gray lines show the reference line (y=0). OLS: Ordinary least squares regression; Lasso: Least absolute shrinkage and selection operator; SVR: Support vector regression; RVR: Relevance vector regression.

## Discussion

In this study, we used structural MRI data to compare the performance of six commonly used linear machine learning algorithms in brain-age prediction and brainPAD in schizophrenia. In patients with schizophrenia, our comparative evaluations showed that ridge regression, Lasso regression, elastic-net regression, SVR, and RVR performed very similarly and yielded comparable results; by contrast, OLS regression performed markedly worse than all other algorithms. BrainPAD has higher in patients with schizophrenia than in healthy individuals although the years of brain-age acceleration varied by algorithm.

We focused specifically on brain structural features derived from FreeSurfer as these are widely used metric examining case-control differences in schizophrenia.^3, 4^ First, using the data from healthy individuals, we demonstrated that the structural feature set entered into the model had a profound effect for the performance of all machine learning algorithms. Incorporating information from cortical thickness, cortical surface area, subcortical volume, and intracranial volume in the same model improved the brain-age prediction in all algorithms. These results reinforce prior studies which found that brain-age can be predicted with a higher accuracy when combining cortical and subcortical brain measures^23, 47^ and suggest that the predictive performance is critically dependent on the input feature data for all machine learning algorithms.^23^ Ridge regression, Lasso regression, and elastic-net regression showed great similarity amongst them in predictive performance as did the SVR and RVR between them; OLS regression generally underperformed and showed a lower similarity with the results of all other algorithms. These findings provide important reference information that can be used in choosing an appropriate machine learning regression algorithm in bran-age prediction in healthy and clinical populations.

In patients with schizophrenia, our comparative evaluations revealed that ridge regression, Lasso regression, elastic-net regression, SVR, and RVR performed very similarly. In contrast, OLS regression performed markedly worse than the other algorithms. Accordingly, significant differences in brainPAD between patients with schizophrenia and healthy individuals were noted for all algorithms except OLS regression. OLS regression is widely used in the neuroscience research as it is relatively easy to implement and can be run quickly with a large number of predictor variables. However, the presence of collinearity among the predictor variables may be a particular vulnerability of OLS regression when applied to brain structural data as there are no regularization techniques within the algorithm. In contrast, both ridge regression and SVR apply L2-norm regularization, Lasso regression includes L1-norm regularization, elastic-net regression includes both L1-norm and L2-norm regularization, and RVR applies regularization through a Gaussian prior. We therefore deduce that the poor performance of OLS regression in this study is likely to be due to the lack of regularization (or penalty) term in the OLS model. The five regularized models provided the similar results in terms of individual brain-age prediction.

The brain regional regression weights in each algorithm reflect the importance of each morphometric feature in brain-age prediction. In patients with schizophrenia, the brain regional regression weights exhibited relatively high inter-algorithm similarity amongst all algorithms, being highest between ridge regression, Lasso regression, elastic-net regression, and RVR. By contrast, the brain regional regression weights derived from OLS regression had the lowest similarity with all the weights from all other algorithms. These findings indicate that the degree of similarity of the brain regions as contributing features varied depending on the regression algorithms which incorporate highly-parameterized models with the heterogeneous brain measures to predict the brain-age. It is worth noting that interpreting the weights of the multivariate brain patterns in the prediction is challenging,^44^ since the prediction is underlined by a combination of several structural features, and that the regional feature weights may not directly provide any neurobiological relevant information. Nevertheless, the present study shows that the similarities of the regularized algorithms extend beyond overall brain-age prediction to include similarity in the regression weights of the individual features.^48^

The mean brainPAD values in patients with schizophrenia over all six algorithms ranged from 3.4 to 5.2 years and were higher than those of healthy individuals (range = −0.49-0.19 years). These findings are aligned with the range of the brainPAD scores in patients with schizophrenia reported in previous studies (range = 2.6-8 years).^16, 18-21^ However, our finding underscore the importance of algorithm selection given that variation in brainPAD was observed despite all analyses being undertaken on the same dataset.

It is also important to acknowledge several limitations that could be addressed in future studies. The key issue is that brain-age prediction can be influenced by factors other than the choice of algorithm such as sample size and the demographic and clinical features of the samples. BrainPAD in patients was associated with the severity for positive and disorganized symptoms suggesting that brain-age prediction in schizophrenia may be influenced by disease severity. Schnack et al have previously shown that the greater deviance in brainPAD is likely to occur within the first five years from disease onset.^16^ Having a single sample allowed to address specifically the effect of the brain-age predictive algorithms but not variation in sample composition. Although we did not test the effect of sample size on the predictive performance of the algorithms, we note that the range of effect sizes found here (*d* range: 0.43-0.66) for each algorithm was similar to that reported by Kaufmann and colleagues (*d* = 0.51) in a much larger sample of over 1000 patients.^17^ Schizophrenia is associated with abnormalities in other structural phenotypes, such as gyrification, as well as in patterns of brain activation and functional connectivity which were not examined here. The present study tested only linear regression models to evaluate the performance of each algorithm for brain-age prediction. Future work should include nonlinear regression models such as Gaussian process regression,^49^ random forest regression,^21^ and deep learning models,^49-51^ but at the cost of interpretability, model complexity and computation time. Theoretically, SVR and RVR can be a nonlinear regression model by applying a nonlinear kernel,^40^ which could be evaluated in future studies as well.

In summary, we provide evidence that linear machine learning algorithms, with the exception of OLS regression, provided similar predictive performance for age prediction on the basis of a combination of cortical and subcortical structural measures. Even when all other parameters were the same, there was variation in the mean brainPAD across algorithms, and therefore algorithm choice is an important source of inter-study variabilitys. Further studies are needed to address variations in brain-age prediction in schizophrenia attributable to other parameters and in other neuroimaging phenotypes. Compared to the other algorithms, OLS regression is the simplest and fastest algorithm, but underperformed in brain-age prediction; therefore, we suggest that the OLS regression should be applied with caution in estimating the brainPAD score in future studies. The Lasso regression performed comparably with other algorithms, but its L1-norm penalty favors sparsity and tends to shrink most weights (coefficients) to zero, which results in a very sparse predictive model based on only a few brain regions. This leads to a sparse solution and provides only selective regional features in a prediction, and thus it should be carefully considered with regard to brain-age acceleration/deceleration in patients in only those few brain regions. RVR introduced as a Bayesian alternative to SVR provided the similar predictive performance and could reduce the model complexity and computational cost. Thus, we would recommend RVR over SVR for brain-age prediction using structural features. Our results suggest that ridge regression is nearly as good as other algorithms, simple to implement and less time-consuming technique than the other regularized algorithms. Collectively, we suggest that researchers carefully report their choices of algorithm used for brain-age prediction to enhance comparability of results across studies.

